# New Approaches for Quantitative Reconstruction of Radiation Dose in Human Blood Cells

**DOI:** 10.1101/640052

**Authors:** Shanaz A. Ghandhi, Igor Shuryak, Shad R. Morton, Sally A. Amundson, David J. Brenner

**Author notes:** Corresponding author: Shanaz A. Ghandhi. Both authors contributed equally to this work and should be considered co-first authors (Shanaz A. Ghandhi, Igor Shuryak). **SAG:**, **IS:**, **SRM:**, **SAA:**, **DJB**.

## Abstract

In the event of a nuclear attack or radiation event, there would be an urgent need for assessing and reconstructing the dose to which hundreds or thousands of individuals were exposed. These measurements would need a rapid assay to facilitate triage and medical management for individuals based on dose. Our approaches to development of rapid assays for reconstructing dose, using transcriptomics, have led to identification of gene sets that have potential to be used in the field; but need further testing. This was a proof-of-principle study for new methods using radiation-responsive genes to generate quantitative, rather than categorical, radiation-dose reconstructions based on a blood sample. We used a new normalization method to reduce effects of variability of gene signals in unirradiated samples across studies; developed a quantitative dose-reconstruction method that is generally under-utilized compared to categorical methods; and combined these to determine a gene-set as a reconstructor. Our dose-reconstruction biomarker was trained on two data sets and tested on two independent ones. It was able to predict dose up to 4.5 Gy with root mean squared error (RMSE) of ± 0.35 Gy on test datasets (same platform), and up to 6.0 Gy with RMSE of 1.74 Gy on another (different platform).

---

The main concern to human health in the event of a nuclear attack or radiation event would be an urgent need for assessing and reconstructing the dose to which hundreds or thousands of individuals were exposed ^1–3^. Many methods are currently being tested for radiation biodosimetry, including well-known methods based on chromosomal damage, that may take days to complete in a high-throughput manner ^4,5^; but also methods that can help to make timely rapid decisions about exposure/dose such as gene expression that have time-to-result within hours. The gold standards used for radiation biodosimetry are cytogenetic assays based on chromosomal changes after DNA damage (reviewed in Sullivan et al.^4^). However, gene expression is an easily measured quantifiable molecular assay, requiring minimal input sample amounts, which can be subjected to multiplexing and modified for high-throughput, and is one of the best candidate molecular measurements for radiation biodosimetry and also have the potential to be used as indicator/predictors of injury. Our approaches to development of these rapid assays for reconstructing dose, using transcriptomics, have led to the identification of gene sets that have the potential to be used in the field; but need to be further tested and subjected to statistical analyses first, to generate a biomarker. A gene signature/biomarker in its basic form will reconstruct the simplest exposure, which is from acute dose rate.

Many studies have been performed using transcriptomics to identify candidates for biodosimetry. These are summarized in the review by Lacombe et al. ^6^, in which a systematic review of available datasets identified genes that could be used for classifying samples by doses above or below 2 Gy. Other studies that have used a comparative approach to identify genes that can reconstruct dose are by Lu et al.^7^ and Macaeva et al.^8^. The goal of the study presented here was to address the simplest type of exposure in humans, representing a total body exposure with an acute dose rate (~1Gy/min), and the response of mRNA levels measured at 24 h after exposure. Once this basic reconstructor signature is established to estimate dose in a quantitative manner, increasingly complex scenarios can also be considered. Gene expression holds promise for distinguishing other relevant characteristics of exposure; including dose rate ^9,10^ such as from exposure to fallout ^11^; and partial shielding and the presence of neutrons ^12,13^, which would be relevant to the blast from an improvised nuclear device ^14^. Genes for discrimination of such dose modifying factors could be tested in future biodosimetric signatures.

To our knowledge, this is the first report of a continuous (non-discreet) dose reconstruction gene-signature with stringent testing on independent datasets for radiation biodosimetry. Our data analysis and results showed that there are some novel aspects to our approach. First, we used a new customized normalization procedure to reduce the effects of variability of gene signals in unirradiated control samples across different studies. Second, we performed quantitative dose reconstructions rather than the more commonly used categorical approach. Finally, by combining these two design methods and applying them to independent datasets we obtained a radiation reconstructor gene-signature that can predict dose and which is more specific compared with other tests based on gene expression that have been reported in the literature.

## Materials and Methods

### Microarray data pre-processing and meta-analysis

In this study, we focused our analyses on human blood irradiated ex vivo to generate a human gene signature for dose reconstruction. We used datasets that had been generated in our group ^9,12,15^ in independent studies performed at different times and using similar acute dose-rate exposures to gamma rays or x-rays as well as a study from Lucas et al ^16^, which used a similar gamma-ray dose range. First, we downloaded the datasets from the NCBI GEO database ^17^, details as in Table 1. We collated each Agilent dataset using BRB-ArrayTools ^18^ and our standard import parameters described previously ^9,15^. For the Lucas et al. dataset (which used an Affymetrix platform, [HG-U133A_2] Affymetrix Human Genome U133A 2.0 Array); the default importer in BRB-ArrayTools was used to collate the data from .cel files. BRB-ArrayTools was then used to normalize the data by median array across common platforms, and filter the data to remove genes with more than 20% missing values. The resulting gene sets were then exported from BRB-ArrayTools for further analysis.

**Table 1.** Ex vivo irradiated Human blood gene expression datasets used in this analysis.

| Datasets for gene signature | NCBI-GEO dataset GSE# | Dose range | PMID/Citation |
| --- | --- | --- | --- |
| 1<sup>st</sup> Training set | 8917 | 0.5 to 8.0 Gy | 18572087 / Paul et al <sup>15</sup> |
| 2<sup>nd</sup> training set (normalizer genes) | 90909 | 0.1 to 4.0 Gy | 28140791 / Broustas et al <sup>12</sup> |
| 1<sup>st</sup> test set | 65292 | 0.56 to 4.45 Gy | 25963628 / Ghandhi et al <sup>9</sup> |
| 2<sup>nd</sup> test set | 58613 | 1.5 to 6.0 Gy | 25255453 / Lucas et al <sup>16</sup> |

### Computational methods to determine and test the continuous dose reconstruction signature

#### Data analysis outline

Two ex vivo photon-irradiated human blood data sets were used for radiation-responsive signature generation (training), and two independent data sets (done at different times) were used for signature testing/validation, as shown in Table 1. These data sets, which contain log_2_ transformed gene signal intensities, were imported into R 3.5.1 software ^19^, where all data analysis steps were performed. These steps were as follows:

a. Using the first training data set, identify genes strongly correlated with radiation dose, which we called “signature” genes.
b. Using the second training data set, identify “normalizer” genes that can reduce the effects of variability in signature gene signal intensities in unirradiated samples across different data sets.
c. Test the combined signature and normalizer gene set on each testing data set by generating continuous dose reconstructions and comparing them with real dose values.

Each step of this approach is described in more detail below.

##### a) Identification of radiation-responsive signature genes

Using the first training data set (Paul et al. GSE8917 ^15^, dose range 0 to 8 Gy); we generated a list of genes with positive Spearman’s correlations with radiation dose, with p-values ≤ 0.05 (with Bonferroni correction) for the Spearman’s correlation coefficient. This analysis was performed using the *cor* and *cor.test* commands in R on each gene in the data set ^19^. As an additional test for robustness to the Bonferroni correction of p-values, synthetic noise variables were added to each data set to serve as benchmarks of reconstructor performance ^20,21^. We added 40,000 synthetic noise variables per data set that were drawn from the normal distribution, and 20,000 per data set from the uniform distribution, using the same mean and SD as all real genes combined. The ratio of noise variables to real genes was approximately 3:1. The rationale for this noise injection into the data set is that only those reconstructors (genes in this case) that outperform all noise variables can be regarded as the strongest ones. Therefore, we retained only those genes for further analysis that: (a) had Bonferroni corrected p-values ≤0.05 for the Spearman’s correlation coefficient with dose, and (2) had Spearman’s correlation coefficient with dose values larger than those for any of the synthetic noise variables.

Because samples exposed to different doses came from the same blood donor, we fitted linear mixed-effects models for each of these significantly radiation-responsive “signature” genes to account for correlations of gene signal intensities by blood donor. The model structure contained a common dose response slope for all donors (fixed effect), but intercepts were allowed to vary by donor (random effect). In other words, the dose response was assumed to be in common for a given gene in all blood donors, but the baseline gene value in unirradiated samples was allowed to vary by donor. More complicated models with random components; for both intercepts and slopes, did not converge in many cases on this data set, so they were not used. We retained only those genes with p-values ≤0.05 (with Bonferroni correction) for the dose-response slope parameter.

To further validate the robustness of the identified radiation-responsive gene signature list, we performed repeated k-means clustering on the same data set using the *kmeans* function in R ^19^. There were 50 repeats, with different initial random numbers seeds. The goal was to identify which genes most frequently appear in the top-scoring cluster (the one with the largest Spearman’s correlation coefficient with radiation dose) out of 50 clustering repeats with different initial random number seeds, and to compare these frequencies with those for synthetic noise variables. We varied the average cluster size from 30 genes per cluster to other values (e.g. 10, 50) to assess the sensitivity of the results to this parameter.

We combined the k-means clustering analysis with: (1) randomization of the outcome variable (radiation dose) by random permutation of the sample labels (doses), and (2) a scenario replacing the dose values with a linear function of a manually selected reconstruction gene-signature (a synthetic noise variable). The goal of these procedures was to assess the false positive rate (i.e. when a gene or set of genes is found to be significantly associated with the randomized dose values in scenario 1) and the sensitivity of the analysis (i.e. what effect size is required to detect the true reconstructor in scenario 2 above).

To assess the correlations between the identified radiation-responsive genes, we calculated the Spearman’s correlation matrix of the genes with each other using the commands *cor* and *e* in R ^19^. The matrix showed that, although the magnitudes of gene signal intensities varied, very strong Spearman’s correlations (≥ 0.73 for any gene pair) were found, suggesting that the dose response shapes for all of these genes are very similar (Figure 1). Consequently, treating each gene as a separate reconstructor of dose would result in severe multi-collinearity. To avoid this problem and group the genes together into one robust reconstructor, we calculated median signal values for all of these signature genes. In other words, the reconstructor value in each sample (i.e. at each dose for each blood donor) was the median value of all the radiation-responsive signature genes identified in the analyses described above (Table 2 and details in Supplementary file 1, signature genes).

**Table 2.** Signature genes for continuous dose reconstruction

| <u>Gene symbol</u> |  |  |
| --- | --- | --- |
| *ANKRA2* | *DRAM1* | *RPS27L* |
| *ANXA4* | *GADD45A* | *SESN1* |
| *ARHGEF3* | *GDF15* | *SLC4A11* |
| *ASCC3* | *IL21R* | *SLC7A6* |
| *ASTN2* | *LIG1* | *TNFRSF10B* |
| *BBC3* | *MAMDC4* | *TRIAP1* |
| *HIST1H2BD* | *MAP4K4* | *UROD* |
| *CDKN1A* | *PCNA* | *VWCE* |
| *DDB2* | *PHPT1* | *WIG1* |
| *EI24* | *PLK3* | *XPC* |
| *FBXO22* | *PPM1D* | *ZNF337* |
| *FDXR* | *PTP4A1* | *ZNF541* |
|  | *REV3L* |  |

**Figure 1.**
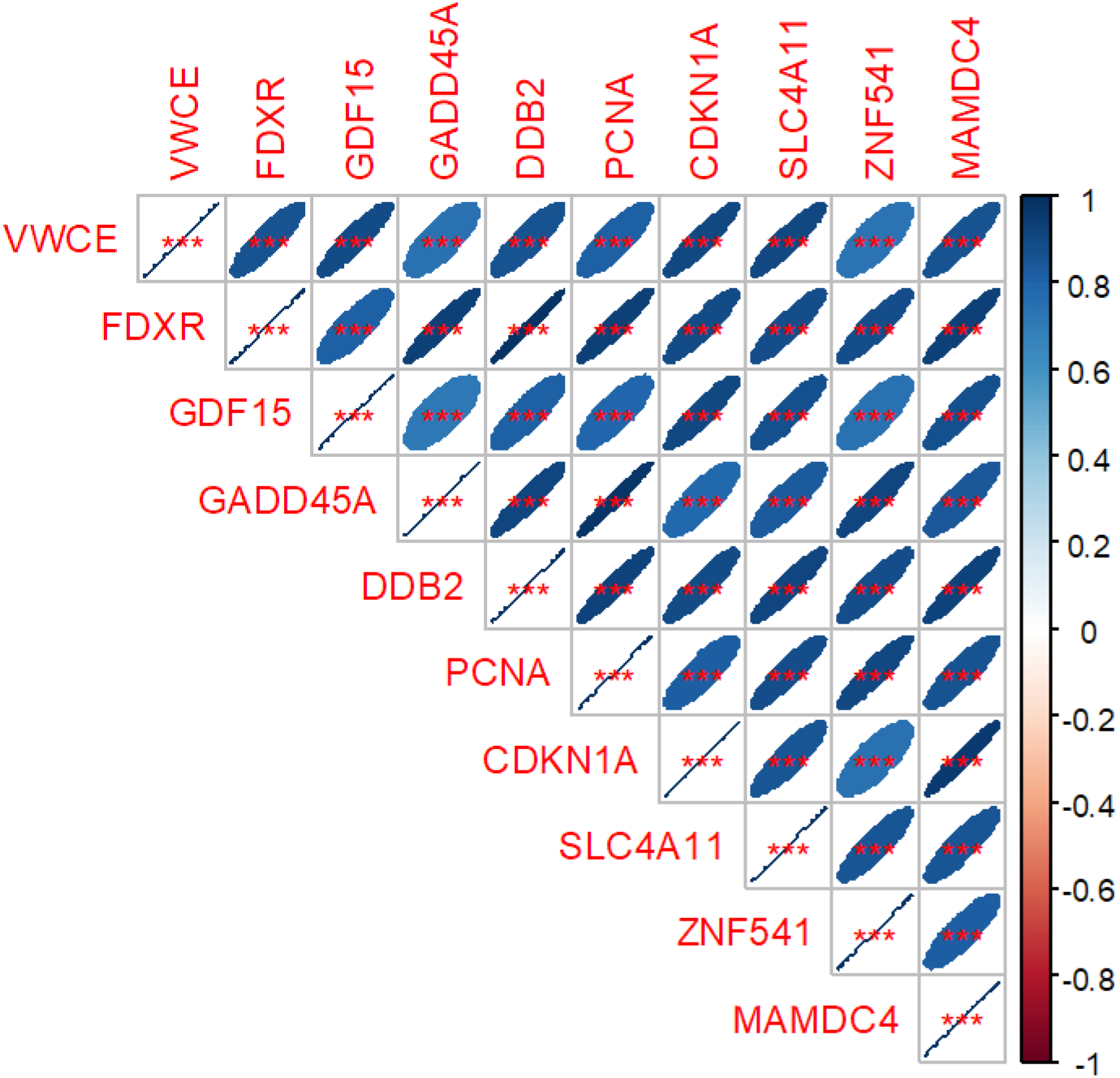
Matrix of Spearman’s correlation coefficients (pairwise, without correction for multiple testing) between 10 selected genes with positive Spearman’s correlations with dose in training data set 1. A color-coded correlation scale is provided on the right of the plot. Crossed out boxes represent meaningless correlations of a given variable with itself. Blue ellipses represent positive correlations, and red ones represent negative correlations. Darker color tones and narrower ellipses represent larger correlation coefficient magnitudes. Red star symbols indicate statistical significance levels: *** indicates p<0.001, ** indicates p<0.01, * indicates p<0.05, no stars indicates p>0.05. These p-values here are intended only for visualization: due to multiple comparisons, only 3 star significance levels are likely to indicate strong associations.

We repeated all the same procedures of gene identification and robustness testing on the same training data set, but looking for genes with negative Spearman’s correlations with radiation dose. However, the correlation coefficient values and subsequent dose reconstruction results for downregulated genes turned out to be much weaker than those for upregulated genes were; so genes with negative Spearman’s correlations with radiation dose were not included in the final analysis (Supplementary file 1, negatively correlated genes).

##### b) Identification of normalizer genes

In our analyses involving independent datasets, performed at different times and from different donors, we have found that baseline expression of genes in unirradiated sample is an issue that can confound the method for dose reconstruction. Although we did not observe any systematic variability in baseline levels of genes, random variability appeared to be considerable (2-4 fold between signals). To reduce the effect of these fluctuations on subsequent dose reconstructions, we developed a method that allowed us to compare gene expression between two independent datasets or platforms using normalizer genes (Supplementary file 1, normalizer genes).

In our analyses, the definition of normalizer gene is one that satisfies the following criteria:

1. the sum of squared differences between a potential normalizer gene’s signal and median signal of the signature gene group has to be as small as possible in unirradiated control samples across ≥2 training data sets.
2. The Spearman’s correlation coefficient for a potential normalizer gene’s signal values with radiation dose across ≥2 training data sets had to be as close to zero as possible.

These criteria were set to identify genes that co-vary with radiation responsive signature genes in unirradiated samples, but do not have radiation responses themselves. We imported the second training data set (Broustas et al. GSE90909 ^12^, 0 to 4 Gy dose range) into R and searched for normalizer genes using both training data sets combined. Two different normalizer gene groups were found, separately for signature genes with positive and negative correlations with radiation dose. We performed normalization across both training data sets by subtracting median signal values for all normalizer genes from median signal values for all signature genes. The retained number of normalizer genes in each group (positive and negative correlations with dose) was selected to be the close to the number of signature genes (Supplementary file 1, normalizer genes).

##### c) Testing the combination of signature + normalizer genes on independent data sets

To create a “standard curve” for relating normalized median gene signals to radiation dose, we fitted a polynomial model (or a robust version that down-weights outliers using the *rlm* function in R ^19^ on data from the two training data sets combined, using dose as the dependent variable and normalized median gene signals as reconstructors (the independent variables). These models had the following structure, where *k*_0_, *k*_1_ and *k*_2_ are adjustable parameters, and *S* is the normalized median gene signal in each sample:

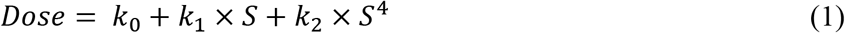

This structure was selected from several alternatives (e.g. different powers of *S* and different numbers of adjustable parameters) using the Akaike information criterion with sample size correction (AICc).

We tested the ability of these fits to reconstruct radiation dose on two independent testing data sets generated using either the same microarray platform (Ghandhi et al. ^12^ with dose range 0 to 4.45 Gy, using acute dose samples only) or a different platform (Lucas et al. ^16^, with dose range 0 to 6 Gy). This was done in each data set by calculating the normalized median gene signal values (*S*) for the signature and normalizer gene groups for each sample. These values were then used as input for the polynomial or robust polynomial models (with original parameter values *k*_0_, *k*_1_ and *k*_2_ derived from fitting training data) to produce dose reconstructions from the testing data. Model performance was evaluated by calculating the coefficient of determination (R^2^) and root mean squared error (RMSE), comparing true and estimated doses.

### Pathway and network analysis and comparisons

We performed gene ontology analysis using DAVID Functional Annotation Tool, ver 6.8 ^22^. We uploaded the human signature gene list to the program and performed functional annotation using biological processes 5, which are ontology-tree child terms. We also uploaded the lists of human signature genes to Ingenuity Pathway Analysis® Software (IPA from Ingenuity®: http://www.ingenuity.com) and performed prediction analysis for upstream regulators. This method identifies potential upstream regulators and ranks them by statistical significance, and provides information about the direction of activation of that regulatory protein based on the downstream gene targets from the gene list. The program provides a z score derived from the number of target genes, their relative expression, and the type of relationship between the regulator and target genes (either activation or inhibition) from the published literature. We compared gene lists using Venny ^23^.

## Results

### Gene Expression results

Published datasets of whole genome gene expression 24 h after ex vivo exposure of peripheral blood from healthy human donors to x-rays or γ-rays (Table 1) were used for this analysis. Genes with more than 20% missing values within a dataset were filtered out and excluded from further analysis. Replicate probes and gene symbols were averaged, which reduced the number of genes in the datasets by ~10% and yielded an average of ~20,000 genes for further analysis from each dataset.

### Radiation responsive gene signature identification and normalization

In the first training data set, we identified 37 genes (Supplementary file 1, signature genes) with strong positive Spearman’s correlations with dose, which withstood our tests for significance and robustness, described in Materials and Methods. Three genes with strong negative correlations with dose were also identified (Supplementary file 1, negatively correlated genes). The genes in each of these two groups were strongly correlated with each other (Figure 1, showing positively correlated genes and Supplementary file 1, negatively correlated genes). In other words, although their signal levels varied, the dose response shapes were very similar for all genes within each group: a nonlinear (concave) increase with dose for the first group, and a nonlinear decrease with dose in the second.

Very similar gene lists were generated by the repeated k-means clustering procedure (described in Materials and Methods). For example, the well-known radiation responsive genes *FDXR* and *DDB2* were found in the top-scoring cluster in 40 out of 50 repeats, whereas several other genes, such as *GADD45A* and *PCNA* were found in 9-10 out of 50 repeats. By comparison, synthetic noise variables were found in the top-scoring cluster in only ≤7 out of 50 repeats. Randomization of the outcome (permutation of dose labels) did not produce any false positives: the highest score for any variable was 11 out of 50, and synthetic noise variables were intermingled with real genes in terms of scores. The alternative test of introducing an artificial dependence of the outcome (dose) on a selected noise variable also performed well in identifying the true predictor in 35 out of 50 repeats. These results demonstrates the ability of proposed methods to separate strong predictors from weak ones and to validate radiation-responsive biomarker signatures generated by previous analyses.

To reduce the effects of variations in baseline levels of signature genes on dose reconstruction, we implemented a search for normalizer genes that co-varied with signature genes in unirradiated samples, but did not have a radiation dose response. This was done by pooling two training data sets. When median signal intensities for normalizer genes were subtracted from the median signal intensities for signature genes, with positive correlations with dose, the results showed a very strong Spearman’s correlation with dose across both training data sets (R^2^ = 0.969, Figure 2). In contrast, the same procedure applied to genes with negative correlations with dose produced much weaker results (Spearman’s correlation with dose was −0.715), and we excluded this gene group from further analysis.

**Figure 2.**
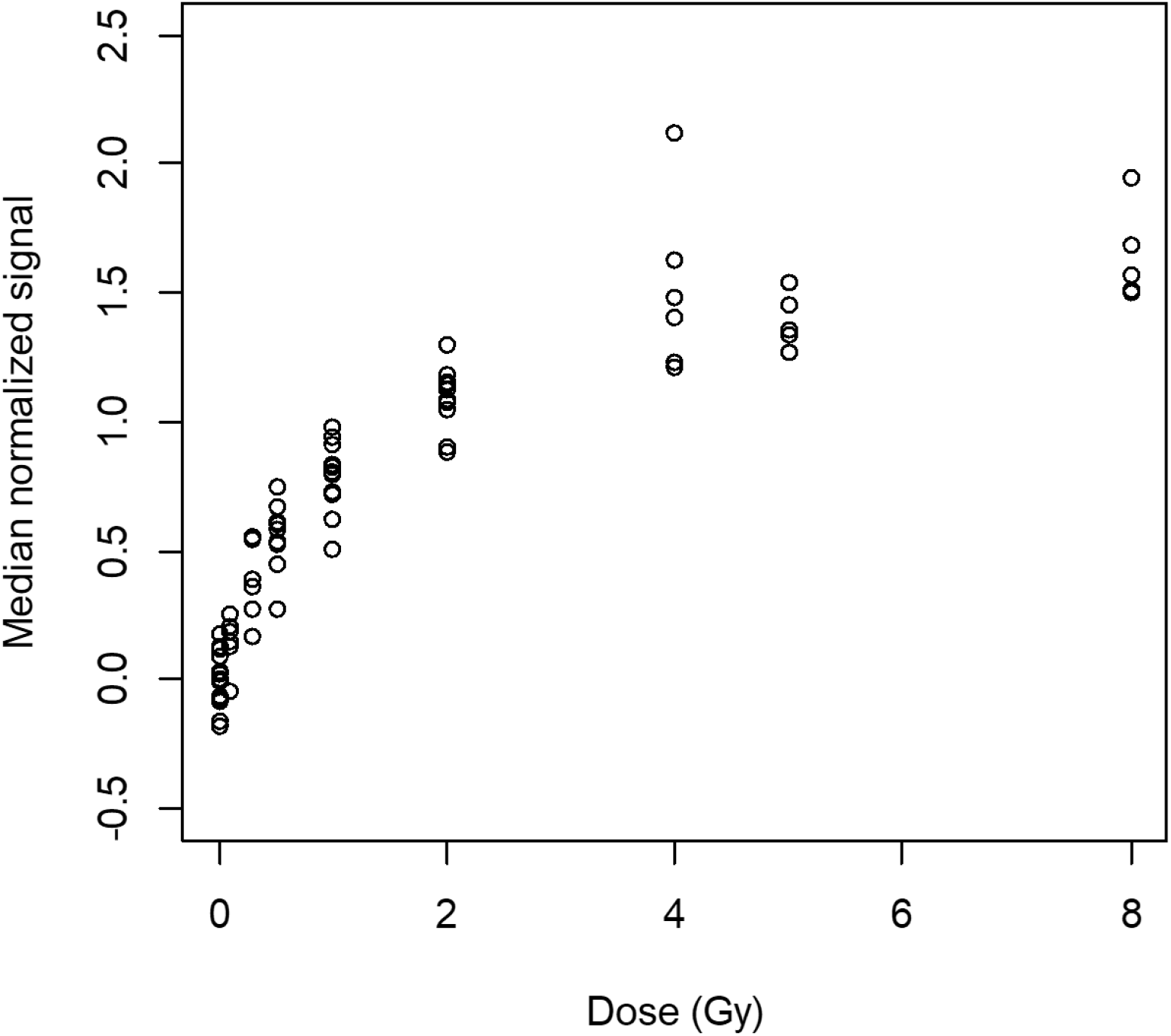
Microarray median normalized gene expression values of the human gene signature in both training data sets combined. Up-regulated genes that were identified using a biostatistics correlation approach in the training data sets are shown in this plot, with corresponding normalizer gene values subtracted. Median log2-transformed signal values are plotted against dose (Gy). Each dot in the graph is a single sample. Spearman’s correlation coefficient with dose was 0.969.

We fitted the polynomial model (Eq. 1 above), using the data for median-normalized signal intensities for signature genes with positive correlations with dose (S) as independent variables, and dose as the dependent variable. Best-fit parameters for the polynomial model were *k*_0_ = −0.43 (standard error = 0.23), *k*_1_ = 2.63 (0.36), *k*_2_ = 0.16 (0.06). For the robust polynomial model, the best-fit parameter values were *k*_0_ = −0.10 (0.08), *k*_1_ = 1.36 (0.13), *k*_2_ = 0.45 (0.02).

### Testing the reconstructor gene signature

Using normalized median gene signal values in the first testing data set (Ghandhi et al. ^9^) as inputs (*S* values) for the polynomial and robust polynomial models, we generated dose reconstructions. The same procedure was performed using second testing data set (Lucas et al. ^16^). Comparisons of true and reconstructed dose values are shown in Table 3 and Figure 3. The dose reconstruction results were very close to the true doses for the first testing data set, which was generated on the same microarray platform as the two training data sets. The results on the second testing data set, which came from a different platform, showed larger error magnitudes. However, even in this case the dose reconstructions were relatively accurate at the highest tested doses.

**Table 3.** Comparison of true doses with reconstructed dose values. SD are standard deviations.

| Testing data set | True dose (Gy) | Reconstructed dose (Gy) using polynomial model |  | Reconstructed dose (Gy) using robust polynomial model |  |
| --- | --- | --- | --- | --- | --- |
|  |  | mean | SD | mean | SD |
| Broustas et al. | 0.00 | -0.07 | 0.30 | 0.09 | 0.16 |
|  | 0.56 | 1.32 | 0.32 | 0.88 | 0.21 |
|  | 2.20 | 2.59 | 0.55 | 1.98 | 0.58 |
|  | 4.40 | 4.18 | 0.13 | 3.96 | 0.19 |
| Lucas et al. | 0.00 | 1.20 | 0.40 | 0.81 | 0.27 |
|  | 1.50 | 4.11 | 0.45 | 3.89 | 0.69 |
|  | 3.00 | 5.16 | 0.84 | 5.60 | 1.40 |
|  | 6.00 | 5.49 | 0.70 | 6.14 | 1.18 |

**Figure 3.**
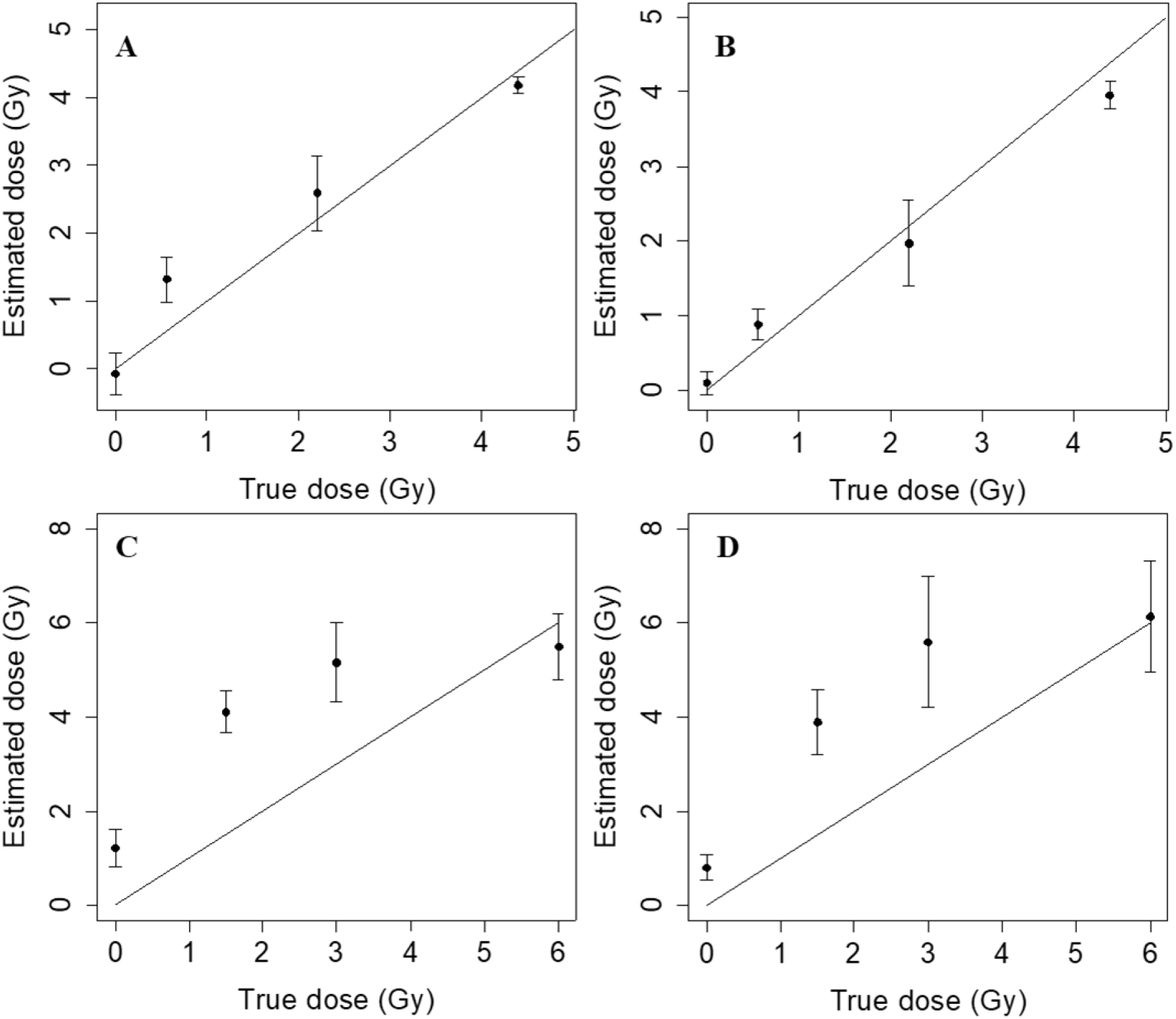
Comparisons of real and estimated radiation doses. **Panel A** Regular polynomial model on testing data set 1. R^2^ = 0.914, RMSE = 0.43 Gy. **Panel B** Robust polynomial model on testing data set 1. R^2^ = 0.952, RMSE = 0.35 Gy. **Panel C** Regular polynomial model on testing data set 2. R^2^ = 0.660, RMSE = 1.74 Gy. **Panel D** Robust polynomial model on testing data set 2. R^2^ = 0.677, RMSE = 1.74 Gy. Error bars represent standard deviations, and the black line the regression line.

### Biological functions of signature genes (networks and pathways, regulators)

We examined the biological significance of the genes in the signature using gene ontology (GO) and pathway analyses. GO analysis using the DAVID database ^22^ suggested enrichment of biological processes in DNA damage response and mitotic cell cycle (Supplementary file 2, worksheet DAVID GO). Apoptosis and cell cycle arrest as well as UV activation of cells were also significantly over-represented among these genes (Benjamini p-value <0.05) and also using IPA network core analysis and biological functions results (Supplementary file 2, worksheet IPA functions). Selecting the pathway category in the ontology and pathway tools, resulted in similar processes being implicated, such as cell death, double strand break repair, mismatch repair and damage of lymphoid cells. We then used IPA to build networks and determine if there were common upstream regulators for this set of genes (Figure 4). Top reconstructed upstream regulators for these genes were AURKB (z score +9.8), ATM (z score 8.7), p38MAPK (z score +6.8), p53 (z score +14.2) and p63 (z score +9.2) (Supplementary file 2, worksheet IPA upstream regulators).

**Figure 4.**
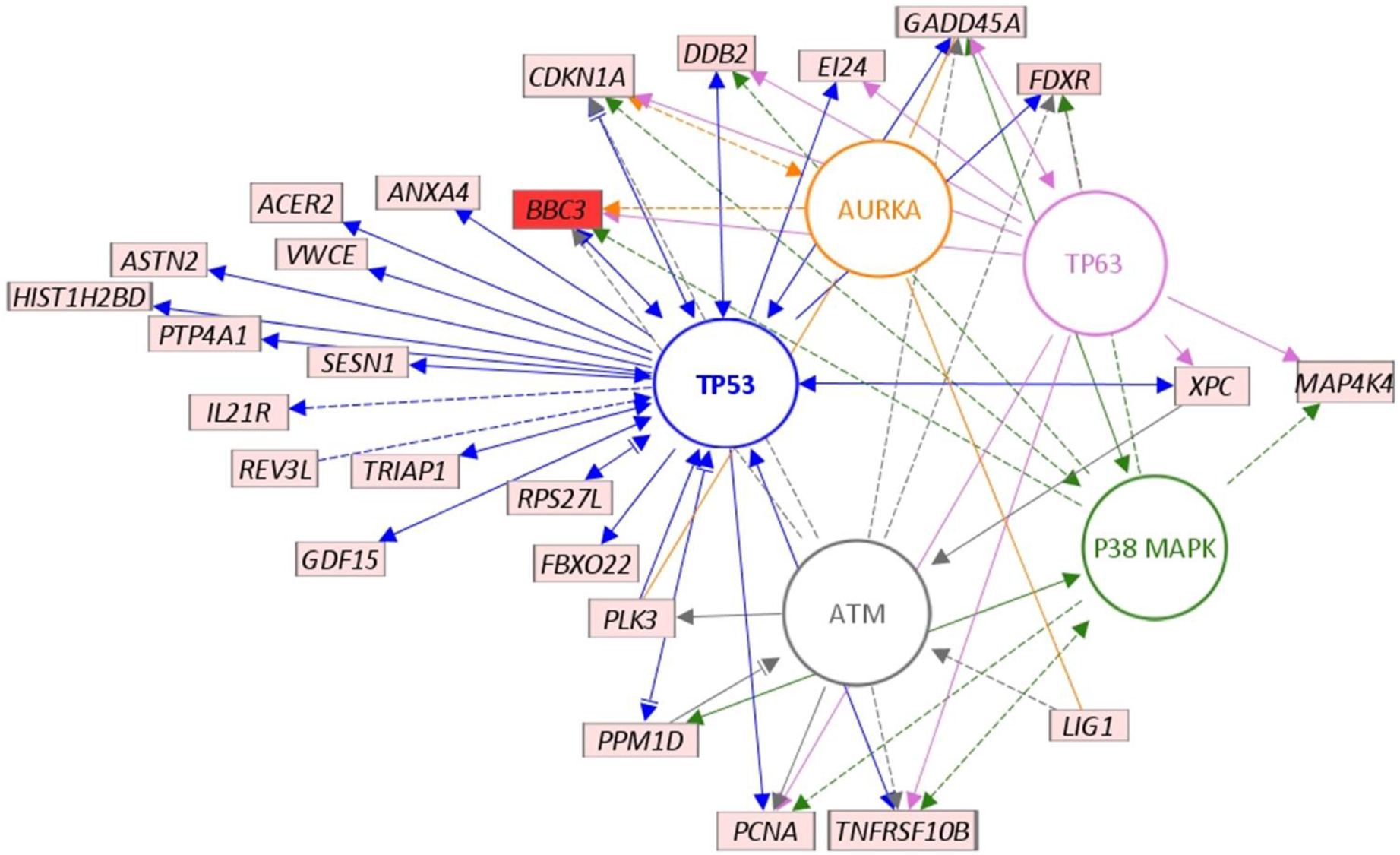
Network analysis showing interactions between the dose-reconstruction signature genes and potential upstream regulators. The network shown here was generated in the Path Designer tool of Ingenuity Pathway Analysis (IPA) program. A subset of signature genes are shown here with connections (indirect, dashed lines; and direct, solid lines/arrows) to the top regulators reconstructed using the IPA algorithm. Nodes/entities are either proteins (circles) or mRNA/genes (rectangles). ATM, AURKA, p38MAPK, TP53 and TP63 were the top regulators (different colors to differentiate between them, with arrows/lines of the same colors connecting the regulator and gene) here with high z scores for activation. All mRNA for signature genes (all up regulated by radiation) shown overlaid with signal/expression values (light pink to bright red).

### Comparison of gene response with other studies

We further visually compared our radiation dose signature genes and their expression across the datasets used in this study in the training/testing analysis. We compared side-by-side gene expression with another dataset (GSE102971, dose range from 0 to 7 Gy) also generated in our group, but performed independently to test similarity of the gene expression response to ex vivo irradiation with non-human primates ^24^. The heat map in Figure 5, displays the scale and changes of signals for 27 of the genes from the reconstructor gene signature (some genes were trimmed for this visualization because of missing values in one or more dataset). GSE102971 data were from different human donors, but were induced similarly for the most of the genes as in GSE8917, which was a training set used here. Some genes such as *CDKN1A* and *ASTN2* were also induced but the mean signal intensities were lower than in the training set. Some genes such as *DDB2* and *TNFRSF10B* showed very similar levels of gene induction and baseline gene expression. We also included neutron data from GSE90909 ^12^, also from our group, as we have found similar responses after neutron doses (last dataset in Figure 5).

**Figure 5.**
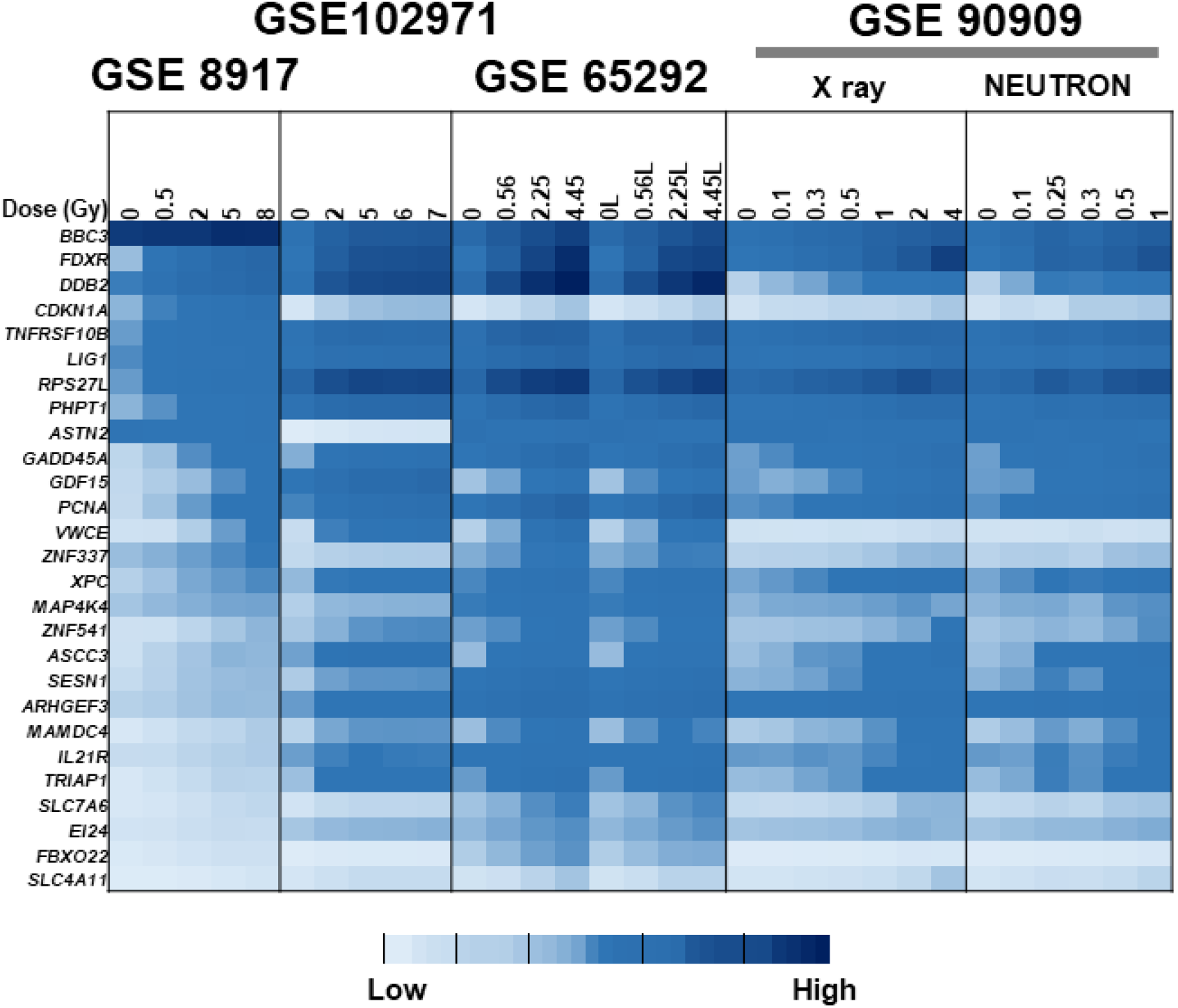
Heat map of gene expression of genes from human blood ex vivo irradiated studies, including an independent study for comparison. Shown here are the expression values for a subset of signature genes (except those that had missing values in some of the datasets) of the sham-irradiated samples (0 Gy) and irradiated samples from different datasets. Starting from the left, first, the training study GSE8917; second, an independent human blood ex vivo irradiation data set GSE102971; third, the first test dataset GSE65292, in which the same donor blood samples were split to study dose-rate effects (suffix L, low dose-rate); and finally, the second training dataset GSE90909 in which the same donor blood samples were split to study LET effects using pure neutrons.

## Discussion

We have developed a robust gene expression signature, which can work effectively to estimate dose in the simplest context. With very stringent biostatistics approaches and by asking a focused question, we have generated a list of target genes that can accurately reconstruct the dose. These results showed that the composition of radiation-responsive signature genes positively correlated with dose is robust against several statistical testing approaches: correlation analysis, mixed effects modeling, synthetic noise.

The accuracy of our human gene signature was in a very good range, especially on a data set from the same platform as the training data (Table 3, Figure 3). Compared with other studies using gene expression results where error for dose reconstruction was within the range ±2.2 Gy ^25^ and with reported microarray-based mean absolute differences of 1.5 to 2.4 Gy for reconstruction of a 4 Gy dose ^26^, our estimation of dose was preferable.

Lacombe et al ^6^ performed a systematic review of 24 independent studies to identify 30 dose reconstruction genes at any time from 2 to 48 h after exposure. They demonstrated the ability of genes from this set to discriminate between doses above or below 2 Gy, but did not test actual dose reconstruction on independent datasets. In contrast, our study focuses on the response at 24 h after exposure, which is the earliest time after a large-scale event when it is thought that first responders may reasonably be able to start assessing the affected population. There were 15 genes in common between the gene-sets reported in our study (Human sig (37)) and Lacombe (31), shown in Figure 6. We also confirmed that most of the genes in the signature identified here were part of the consensus gene set (Paul (64) in Figure 6) used to classify samples by dose in the initial analysis of the study that provided our first training dataset ^15^. The genes common to all three signatures were *ASCC3*, *BBC3*, *CDKN1A*, *DDB2*, *EI24*, *FBXO22*, *FDXR*, *GADD45A*, *PCNA*, *PHPT1*, *RPS27L*, *SESN1*, *TNFRSF10B*, *TRIAP1* and *XPC* all of which are known-radiation genes ^27–32^. There were an additional 16 genes in common between our robust dose reconstructor and the Paul (64) consensus gene set (*ANKRA2*, *ANXA4*, *ARHGEF3*, *ASTN2*, *GDF15*, *IL21R*, *LIG1*, *MAMDC4*, *PLK3*, *PPM1D*, *PTP4A1*, *SLC4A11*, *SLC7A6*, *UROD*, *VWCE* and *ZNF337*). Interestingly, some genes (*HIST1H2BD*, *DRAM1*, *MAP4K4*, *REV3L*, *WIG1* and *ZNF541*) were only included in the signature identified in this study; most of these are involved in the p53 and/or radiation response ^33–38^.

**Figure 6.**
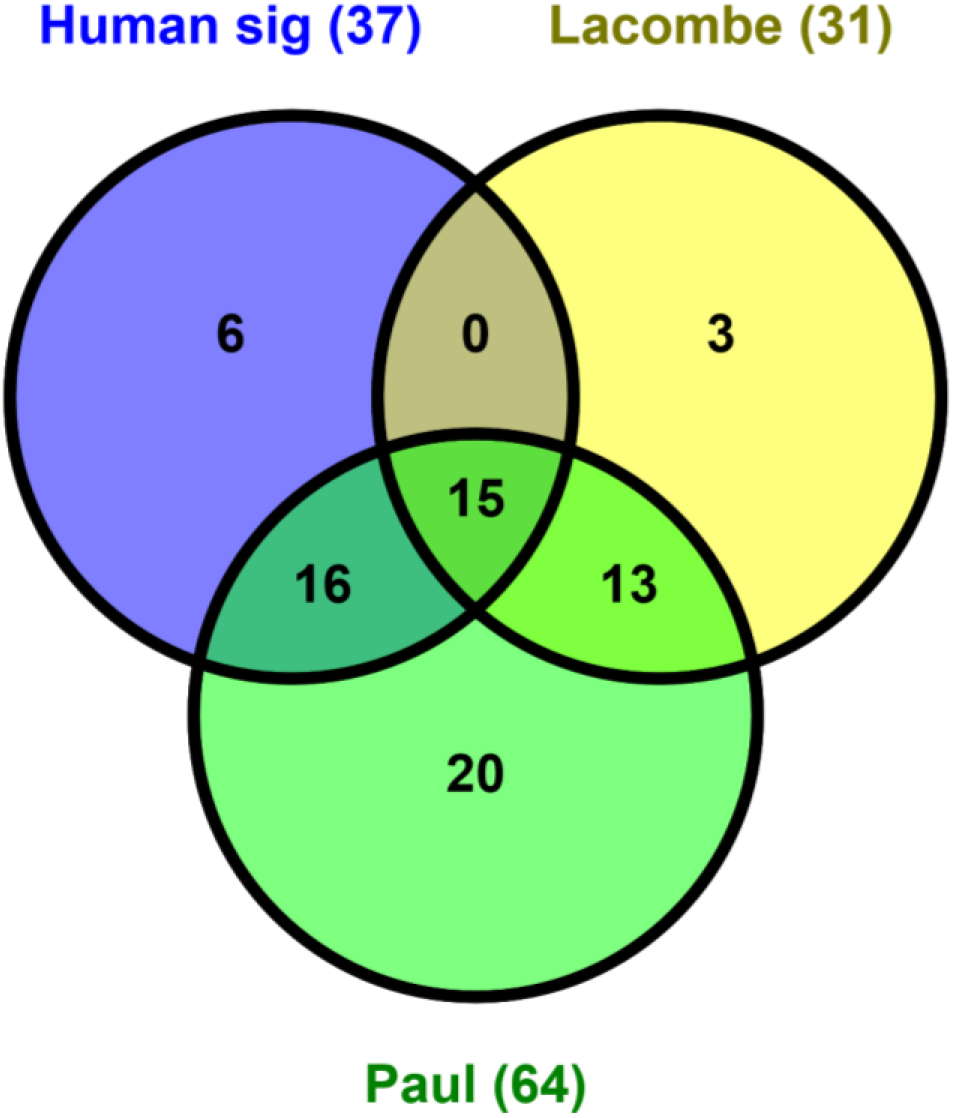
Venn diagram showing overlap between our signature genes and others. We compared overlap between gene signatures published in two other studies. The Human sig (signature) genes were the 37 genes from the biostatistics method described here, the Lacombe study also used a meta-analysis approach from which 31 genes were identified from published studies at different times post-irradiation, and the Paul et al, 64 consensus-gene set that has been used in numerous meta-analysis studies of radiation biomarker identification.

The ultimate goal of radiation biodosimetry is to reconstruct the dose to people who were exposed in vivo. Several studies have reported that some, but not all, of the in vivo gene expression response to radiation measured in blood cells is recapitulated when the blood is irradiated ex vivo, and that informed selection of genes allows signatures based on ex vivo exposures to reconstruct dose levels of in vivo exposures ^16,39–41^. Many of the genes commonly suggested for biodosimetry have also been shown to respond to the stress of blood cells being cultured outside the body ^39^. Implementation of a dose-reconstruction gene signature may require modification to account for the in vivo context.

The goal of the analysis described here was to determine a core gene-signature panel to reconstruct the gamma-equivalent dose in irradiated human blood in an experimental scenario that simulates the potential real life exposures that may occur after a radiation accident or bomb. The set of genes selected by these analyses are mostly well-known radiation response genes that correlate very well with dose. We also showed as a proof-of-principle that a strategy to find normalizer genes to correct for variability in these signature genes in unirradiated control samples from donors across datasets, could work to generate continuous dose reconstructions in a training/testing framework.

## Conclusions

We identified radiation-responsive “signature” genes with continuous dose responses and strong positive correlations with dose that are consistent across several data sets. The gene group with negative correlations with dose was much smaller/weaker in these ex vivo blood data sets. The radiation-responsive “signature” genes make sense biologically, overlap with previous findings, and stood up to various tests: addition of synthetic noise, mixed effects modeling of donor effects, different forms of clustering. Therefore, we propose our present gene set, that has the added strength of being tested across independent datasets and across more than one platform, will perform well to predict the acute dose to an individual; and in our tests, the most accurate prediction of dose had an acceptable level of error of ± 0.35 Gy.

## Declarations

### Ethics, approval, and consent to participate

No animal or human experiments were performed as part of this study.

### Availability of data and material

Microarray datasets used for meta-analyses in this study are publically available in the NCBI Gene Expression Omnibus database (https://www.ncbi.nlm.nih.gov/geo/) under accession numbers GSE8917, GSE90909, GSE65292, GSE102971 and GSE58613 (independent group).

### Author’s contributions

IS performed all biostatistics analyses and helped to draft the manuscript. SAG and SAA conceived the study, helped with the analyses and drafted the manuscript. DJB reviewed the biostatistics analysis and helped draft the manuscript and SRM helped prepare the manuscript and figures.

## List of Supplementary files

**Supplementary file 1** Signature genes and annotations

**Supplementary file 2** Gene ontology and network analysis

